# Spatiotemporal distribution of Delta-like protein 1 during mouse pituitary ontogeny and its relationship with differentiating endocrine cell populations

**DOI:** 10.64898/2026.06.26.734803

**Authors:** Ricardo Reyes, Alexis Rufino-Gómez, Carmen Díaz, Aixa R Bello

## Abstract

Delta-like protein 1 (DLK1) is a transmembrane protein involved in the regulation of cellular differentiation and stem cell maintenance in several tissues, including the pituitary gland. Although DLK1 expression has been reported in the adult pituitary, its spatiotemporal distribution during mouse pituitary development remains incompletely characterized. The aim of this study was to analyse the distribution of DLK1 during embryonic and postnatal development of the mouse pituitary gland and to characterize its relationship with hormone-producing cell populations. Immunohistochemistry was performed in Swiss albino mice from embryonic day 9.5 (e9.5) to postnatal day 15 (p15). Double immunofluorescence was used at e18.5 and p15 to examine the association of DLK1 immunoreactivity with ACTH-, TSH-, GH-, FSH- and PRL-producing cells.

DLK1 immunoreactivity was detected from the earliest stages of pituitary development in Rathke’s pouch and the ventral diencephalon. During embryonic development, DLK1-ir cells were widely distributed throughout adenohypophyseal and neurohypophyseal primordia. As development progressed, DLK1 immunoreactivity became progressively regionalized within the anterior, intermediate and tuberal lobes, as well as in the median eminence of the hypothalamus and the posterior pituitary lobe. Colocalization of DLK1 with each of the hormone-producing cell populations analysed was observed at e18.5 and p15. Semiquantitative analysis at p15 indicated that approximately 32% of adenohypophyseal cells were DLK1-immunoreactive.

These findings provide a detailed description of the spatiotemporal distribution of DLK1 during mouse pituitary ontogeny and suggest that DLK1 immunoreactivity is associated with differentiating endocrine cell populations throughout pituitary development.

## Introduction

The adenohypophysis, including the anterior (AL), intermediate (IL) and tuberal (TL) lobes of the pituitary gland, derives from an invagination of the primitive oral ectoderm known as Rathke’s pouch, whereas the posterior pituitary (posterior lobe, PL) develops from the neural ectoderm of the ventral diencephalon. The median eminence, a specialized region of the hypothalamus, also arises from the ventral diencephalon and develops in close anatomical association with the posterior pituitary. During pituitary organogenesis, progenitor cells progressively differentiate into specific hormone-producing cell populations through coordinated spatial and temporal developmental events.^1–5^

Pituitary development is regulated by multiple signalling pathways and transcription factors, including BMP, FGF, SHH, WNT and NOTCH signalling systems, which participate in progenitor maintenance, lineage specification and endocrine cell differentiation.^6–13^ In mammals, many of these signalling molecules are produced both extrinsically in the ventral hypothalamus and intrinsically within the developing pituitary gland,^8–13^ while similar spatiotemporal expression patterns have also been described during avian pituitary development.^14^

Pituitary progenitor cells express PROP1, a paired-like homeodomain transcription factor required for pituitary identity and endocrine lineage commitment.^15^ PROP1 activity is associated with activation of Pou1f1 (formerly Pit1), which is involved in somatotroph, lactotroph and thyrotroph differentiation, whereas Tbx19 (formerly Tpit) is required for corticotroph and melanotroph development.^8,9,11^ The NOTCH signalling pathway has also been associated with progenitor maintenance and proliferation during pituitary development, as well as with regulation of Prop1 expression.^11,16–18^ NOTCH receptors are activated by membrane-bound ligands belonging mainly to the DELTA-like and JAGGED families.^19,20^

Several studies have shown that Delta-like protein 1 (DLK1), a non-canonical NOTCH ligand belonging to the epidermal growth factor-like protein family, may modulate NOTCH signalling activity.^21–25^ DLK1 expression has been associated with progenitor or incompletely differentiated cellular states in several developmental systems.^26–28^ In the pituitary gland, DLK1 has been related to growth hormone regulation in cultured pituitary cells,^29^ and DLK1 immunoreactivity has been identified in hormone-producing cells of the adult mouse pituitary.^30^ Likewise, Dlk1 mRNA expression has been reported in the developing adenohypophysis of rodents,^28,31^ and DLK1 protein has been detected in developing rat pituitary endocrine cell populations, particularly in ACTH- and GH-producing cells.^32^

More recently, Scagliotti et al.^33^ analysed DLK1 expression in the embryonic mouse pituitary gland and highlighted the relationship between Dlk1 dosage, stem cell populations and pituitary size regulation. Whereas Scagliotti et al.^33^ focused primarily on stem-cell populations and the effects of Dlk1 dosage on pituitary growth, the spatiotemporal distribution of DLK1 during pituitary regionalization and its association with differentiating hormone-producing cell populations remain incompletely characterized. Therefore, a detailed developmental analysis of DLK1 protein distribution throughout pituitary ontogeny is still lacking.

Therefore, the aim of the present study was to analyse the spatiotemporal distribution of DLK1-immunoreactive cells during mouse pituitary development, with special attention to Rathke’s pouch, adenohypophyseal lobes, the posterior pituitary and the median eminence, and to characterize their relationship with differentiating hormone-producing cell populations.

## Materials and methods

### Animals and tissue collection

All experimental procedures involving animals were conducted in accordance with the European Union Directive 2010/63/EU and Spanish legislation for animal experimentation (Real Decreto 53/2013) and were approved by the Ethics Committee for Animal Experimentation of the University of La Laguna (CEIBA2023-3304). This study was conducted and reported in accordance with the ARRIVE 2.0 guidelines for animal research.

Thirty-six Swiss albino mouse embryos and postnatal animals were analyzed at embryonic days (e) 9.5, 11.5, 13.5, 14.5, 17.5, 18.5 and 19.5, and postnatal days (p) 7 and 15 (n = 4 per developmental stage). The day of vaginal plug detection was considered embryonic day 0.5 (e0.5), according to the conventional staging system for mouse embryonic development.^34^

Embryos at early developmental stages were collected as whole embryos, whereas older embryos and postnatal animals were decapitated prior to fixation. Samples were rinsed in 0.1 M sodium phosphate buffer (pH 7.4) and fixed in 4% paraformaldehyde in phosphate buffer for 48 h at room temperature. After fixation, tissues were dehydrated through graded ethanol series, embedded in Paraplast® (Sigma-Aldrich, St. Louis, MO, USA) and sectioned sagittally at 5 µm using a Shandon Finesse 325 microtome. Sections were mounted on Polysine®-coated slides (Thermo Scientific, Germany).

For general histological examination, sections were stained with haematoxylin and erythrosine.

### Immunohistochemistry

Sections were deparaffinized, rehydrated through decreasing ethanol concentrations and washed in Tris-buffered saline (TBS; 0.05 M Trizma Base, 0.9% NaCl, pH 7.4), which was used in all subsequent washes and incubations.

For immunohistochemical detection of DLK1, sections were preincubated in TBS containing 5% fetal bovine serum (FBS) and 0.2% Triton X-100 to reduce non-specific binding. Sections were then incubated overnight at room temperature with a mouse monoclonal anti-DLK1 antibody (1:1000; Santa Cruz Biotechnology Inc., Germany). After washing, sections were incubated for 1 h at room temperature with biotin-SP goat anti-mouse IgG (1:1000; Jackson ImmunoResearch Laboratories, West Grove, PA, USA), followed by peroxidase (POD)-conjugated streptavidin (1:1000; Jackson ImmunoResearch Laboratories, West Grove, PA, USA). Peroxidase activity was revealed using the glucose oxidase-DAB-nickel method in the presence of 0.01% hydrogen peroxide.^35^

Immunostaining specificity was evaluated in control sections by omitting the primary antibody. Specific non-immune serum (1:500) was used as a negative control for each primary antibodies.

### Double immunofluorescence

To analyse the relationship between DLK1 expression and pituitary hormone-producing cell populations, double immunofluorescence was performed on selected sections.

After deparaffinization and rehydration, sections were incubated in TBS containing 5% FBS and 0.2% Triton X-100 to reduce non-specific binding prior to primary antibody incubation. Sections were then incubated overnight at room temperature with the mouse monoclonal anti-DLK1 antibody together with antibodies against different pituitary hormones.

The following primary antibodies were used: mouse monoclonal anti-DLK1 (1:1000; Santa Cruz Biotechnology Inc., Germany), guinea pig polyclonal anti-PRL (1:500; United States Biological, Swampscott, MA, USA), rabbit polyclonal anti-ACTH (1:1000), rabbit polyclonal anti-TSH (1:600), rabbit polyclonal anti-FSH (1:800), and rabbit polyclonal anti-GH (1:800) (G. Tramu, University of Bordeaux 1, Bordeaux, France, for ACTH, TSH and FSH; Chemicon-Merck-Millipore, Schwalbach, Germany, for GH). The immunological specificity of the ACTH, TSH and FSH antisera has been previously characterized and validated in immunohistochemical studies.^36–39^

Species-specific secondary antibodies were used to minimize cross-reactivity between immunodetection channels. DLK1 immunoreactivity was detected using biotin-SP goat anti-mouse IgG (1:1000; Jackson ImmunoResearch Laboratories, West Grove, PA, USA), followed by Cy3-conjugated streptavidin (1:1000; Jackson ImmunoResearch Laboratories). Pituitary hormones were detected using Alexa Fluor 488-conjugated goat anti-rabbit IgG (1:300; Jackson ImmunoResearch Laboratories) and Alexa Fluor 488-conjugated goat anti-guinea pig IgG (1:300; Jackson ImmunoResearch Laboratories).

Control sections were processed by omitting one or both primary antibodies and, where appropriate, replacing the primary antibody with non-immune rabbit serum (1:500). In addition, adult mouse heart and kidney, reported to lack detectable DLK1 immunoreactivity,^40,41^ were processed using the same immunohistochemical procedure as negative tissue controls (Figure S2).

The characteristics and specificity of all primary and secondary antibodies and detection reagents used in this study are summarized in Table 1.

**Table 1.** Characteristics of the primary and secondary antibodies and detection reagents used in this study.

| Category | Target/reagent | Antibody/reagent | Antigen/specificity | Host species | Source | Cat. No. | Dilution |
| --- | --- | --- | --- | --- | --- | --- | --- |
| Primary antibodies | ACTH | Anti-ACTH (1–24) | Synthetic ACTH (amino acids 1–24) | Rabbit polyclonal | G. Tramu, University of Bordeaux 1, Bordeaux, France | – | 1:1000 |
|  | DLK1 | Anti-DLK1 (B-7) | Human DLK1, amino acids 266–383 | Mouse monoclonal | Santa Cruz Biotechnology Inc., Germany | sc-376755 | 1:1000 |
| | FSH | Anti-hβFSH | Human βFSH | Rabbit polyclonal | G. Tramu, University of Bordeaux 1, Bordeaux, France | – | 1:800 |
|  | GH | Anti-Growth Hormone | Purified growth hormone | Rabbit polyclonal | Chemicon-Merck-Millipore, Schwalbach, Germany | AB940 | 1:800 |
| | PRL | Anti-Prolactin (PRL) Pab Gp × Hu | Purified prolactin hormone | Guinea pig polyclonal | United States Biological, Swampscott, MA, USA | P9009-16 | 1:500 |
| | TSH | Anti-hβTSH | Human βTSH | Rabbit polyclonal | G. Tramu, University of Bordeaux 1, Bordeaux, France | – | 1:600 |
| Secondary antibodies | Mouse IgG | Biotin-SP Goat anti-Mouse IgG | Mouse IgG | Goat | Jackson ImmunoResearch Laboratories, West Grove, PA, USA | 115-066-006 | 1:1000 |
|  | Rabbit IgG | Alexa Fluor 488<br>Goat anti-Rabbit<br>IgG | Rabbit IgG | Goat | Jackson<br>ImmunoResearch<br>Laboratories, West<br>Grove, PA, USA | 111-546-047 | 1:300 |
|  | Guinea pig IgG | Alexa Fluor 488<br>Goat anti-Guinea<br>Pig IgG | Guinea pig IgG | Goat | Jackson<br>ImmunoResearch<br>Laboratories, West<br>Grove, PA, USA | 106-547-008 | 1:300 |
| **Detection**<br>**reagents** | Biotin | POD-conjugated<br>Streptavidin | Biotin | — | Jackson<br>ImmunoResearch<br>Laboratories, West<br>Grove, PA, USA | 016-030-084 | 1:1000 |
|  | Biotin | Cy3-conjugated<br>Streptavidin | Biotin | — | Jackson<br>ImmunoResearch<br>Laboratories, West<br>Grove, PA, USA | 016-160-084 | 1:1000 |
| **Nuclear**<br>**counterstain** | Nuclear DNA | DAPI (4',6-diamidino-2-phenylindole<br>dihydrochloride) | DNA | — | Sigma-Aldrich,<br>Spain | D9542 | 0.5 µg/mL,<br>1 min |

### Image acquisition and analysis

Sections were examined using a Leica DM4000B light microscope equipped with epifluorescence optics and a Leica DFC300 FX digital camera (Leica Microsystems, Wetzlar, Germany). Fluorescence channels were acquired sequentially and merged digitally after image acquisition. For double-immunofluorescence images presented in the main figures, DLK1 and hormone channels were merged to facilitate visualization of DLK1 and hormone colocalization. DAPI counterstaining was additionally examined in selected sections to assess tissue integrity and cellularity (Figure S1). Images were processed using Leica Q-Win® software (Leica Microsystems, Barcelona, Spain) and ImageJ software.

Because DLK1 immunoreactivity frequently displayed diffuse cytoplasmic and membranous labeling patterns, identification of cells showing DLK1 and hormone colocalization was based on conservative morphological criteria.

### Semiquantitative analysis

To obtain a semiquantitative estimation of DLK1-positive cells in the anterior pituitary lobe, four p15 mouse pituitaries were analyzed. Eighteen sagittal sections were obtained from each pituitary gland by mounting three sections per slide on six slides.

Sections were processed for DLK1 immunofluorescence and counterstained with DAPI (0.5 µg/mL; Sigma-Aldrich, Spain) for 1 min to identify cell nuclei while minimizing excessive nuclear fluorescence. Five randomly selected fields per slide were digitized using Leica Q-Win® software. Total cell number (DAPI-positive nuclei) and DLK1-immunoreactive cells were quantified to estimate the relative proportion of DLK1-positive cells within the anterior lobe (Equation 1).

To assess the distribution of DLK1 colocalization with hormone-producing cell populations, double immunofluorescence was performed for DLK1 together with ACTH, TSH, GH, FSH or PRL. DLK1-positive cells and cells showing DLK1 and hormone colocalization were counted in the same randomly selected fields to estimate the relative proportion of DLK1-immunoreactive cells showing colocalization with each hormone-producing cell population (Equation 2). Only cell profiles showing immunostained cytoplasm and a recognizable nuclear profile were included in the analysis. Small stained fragments and artifacts were automatically excluded by applying minimum object size thresholds. Immunoreactivity was evaluated according to fluorescence intensity values above background levels obtained from control sections. Cell counts were independently performed by the first and second authors (RR and ARG) under blinded conditions.

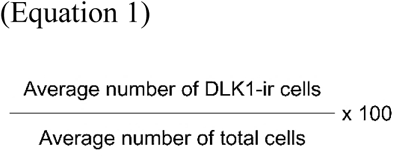

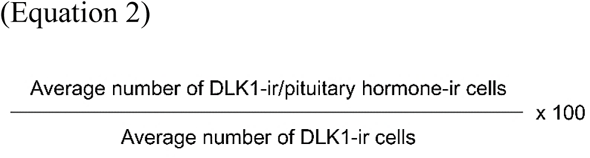

## Results

### Spatiotemporal distribution of DLK1 during pituitary development

At embryonic day 9.5 (e9.5), DLK1 immunoreactivity was already observed in the ventral diencephalic neuroepithelium and in the oral ectoderm associated with the developing Rathke’s pouch (Figure 1a,b). During subsequent developmental stages, DLK1-ir cells progressively increased in both intensity and distribution throughout the developing pituitary gland.

**Figure 1.**
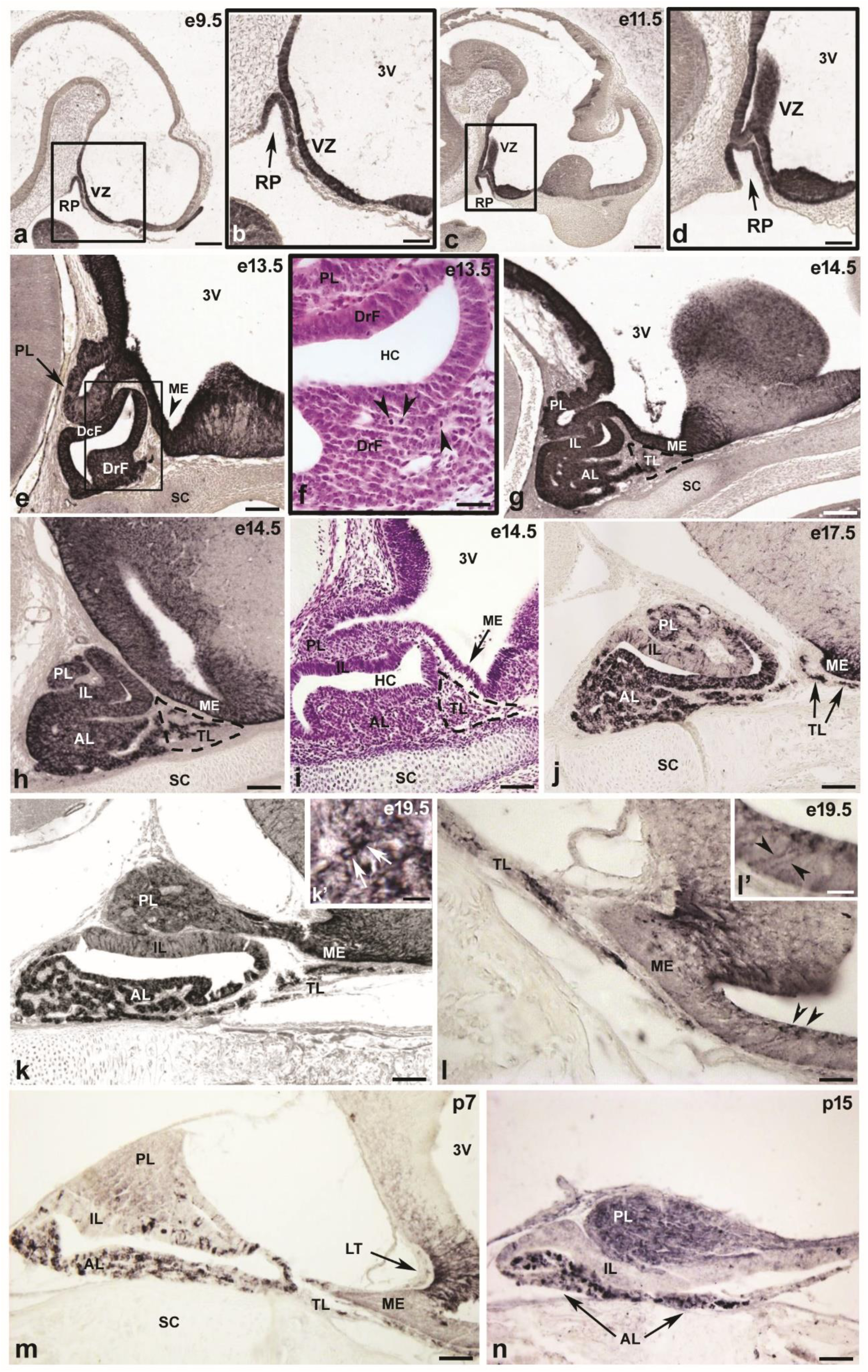
Presence of DLK1 immunoreactivity during pituitary development in sagittal sections of Swiss albino mice. The caudorostral (C, R) and ventrodorsal (V, D) spatial directions are indicated in (a). Rathke’s pouch and ventricular zone of the hypothalamic primordium are DLK1-immunoreactive (ir) in mouse embryos at stages e9.5 (a) and e11.5 (c). Magnified framed areas show immunoreactivity in Rathke’s pouch (arrow) (b and d). Sagittal section of Rathke’s pouch at stage e13.5 exhibiting DLK1 immunoreactivity in both faces of Rathke’s pouch (dorsorostral and dorsocaudal), and developing posterior lobe (arrow) and median eminence (arrowhead) (e). Magnified frame illustrates haematoxylin-erythrosine staining in the dorsorostral face of the pouch showing mitotic profiles (arrowheads) (f). DLK1 immunoreactivity in anterior, intermediate and tuberal lobes (the latter with a dashed outline) in addition to posterior lobe and median eminence of a mouse pituitary in the embryonic stage e14.5 (g and h) and a comparable section level stained with haematoxylin-erythrosine at e14.5 (i). DLK1 immunoreactivity in embryonic mouse pituitary gland at stage e17.5 (j). DLK1 expression at stage e19.5 (k). Posterior lobe magnification (insert) with DLK1-ir pituicytes (white arrows) (k’). Images (l and l’, insert) show tuberal lobe and median eminence at stage e19.5 with DLK1-ir tanycytes (arrowheads). The images (m) and (n) show the DLK1 immunoreactivity in a sagittal section of postnatal pituitary gland in stages p7 and p15 respectively. AL: Anterior lobe; DrF: Dorsorostral face; DcF: Dorsocaudal face; IL: Intermediate lobe; HC: Hypophyseal cleft; ME: Median eminence, PL: Posterior lobe; RP: Rathke’s pouch; TL: Tuberal lobe; VZ, ventricular zone; 3V: Third ventricle. Scale bars = (a,c): 100 µm, (b,d,e,g,h,i,j,k,m,n): 50 µm, (f,k’,l,l’): 20 µm.

At e11.5, DLK1 immunoreactivity was detected in the developing Rathke’s pouch epithelium and in the adjacent ventral diencephalon (Figure 1c,d). At e13.5 and e14.5, DLK1-ir cells were widely distributed throughout the adenohypophyseal primordium, particularly in the developing anterior lobe (AL), intermediate lobe (IL) and tuberal lobe (TL), while immunoreactivity was also observed in the neurohypophyseal region (Figure 1e,g,h). Numerous mitotic profiles were observed within the developing adenohypophyseal tissue during these stages (Figure 1f). The corresponding H-E-stained section is shown in Figure 1i.

No specific DLK1 immunoreactivity was observed in the negative control sections (Figure S2).

At e17.5 and e18.5, DLK1 immunoreactivity became more clearly regionalized within the pituitary gland. In the AL, DLK1-ir cells were broadly distributed throughout the lobe, whereas in the IL, immunoreactive cells were preferentially located adjacent to the developing posterior lobe (PL) and to the hypophysary cleft (HC). In the tuberal lobe (TL), DLK1-ir cells were detected surrounding the infundibular region. DLK1 immunoreactivity was also observed in the median eminence and PL (Figure 1j).

At e19.5, DLK1-ir cells remained abundant in the AL, whereas a heterogeneous distribution pattern became more evident in the IL and TL (Figure 1k,l). In the posterior pituitary, DLK1-immunoreactivity was detected in pituicytes and nerve fibres (Figure 1k, k’). In addition, DLK1-ir fibres and tanycytes were observed in the median eminence (Figure 1k,l, ĺ). In postnatal stages (p7 and p15), DLK1 immunoreactivity persisted throughout the adenohypophysis, posterior pituitary and median eminence (Figure 1m,n). In the AL, DLK1-ir cells displayed heterogeneous staining intensity and distribution, whereas in the IL immunoreactivity decreased in p7 until it almost disappeared at p15 (Figure 1m,n). In the TL and median eminence immunoreactivity was also reduced, with a lower immunoreaction observed in both p7 and p15, while in the PL, lower immunoreactivity was observed in p7 and increased in p15 (Figure 1m,n).

### DLK1 immunoreactivity in hormone-producing cell populations during embryonic development

Minor inter-individual differences in DLK1 immunofluorescence intensity were observed among different embryos of the same developmental stage. Nevertheless, the overall regional distribution pattern of DLK1 immunoreactivity remained highly consistent across specimens.

Double immunofluorescence analyses were performed at e18.5, representing a late embryonic stage at which all major hormone-producing cell populations are undergoing differentiation. No specific fluorescence signal was observed in the corresponding negative control sections (Figure S2).

At e18.5, double-labelled cells for DLK1 and ACTH were observed in the AL (Figure 2a–c). Likewise, double-labelled cells for DLK1 and TSH (Figure 2d–f), GH (Figure 2g–i), FSH (Figure 2j–l) or PRL (Figure 2m–o) were identified within the AL. DLK1 immunoreactivity frequently displayed diffuse cytoplasmic and membranous labeling, whereas hormone immunoreactivity showed more restricted cytoplasmic distribution patterns.

**Figure 2.**
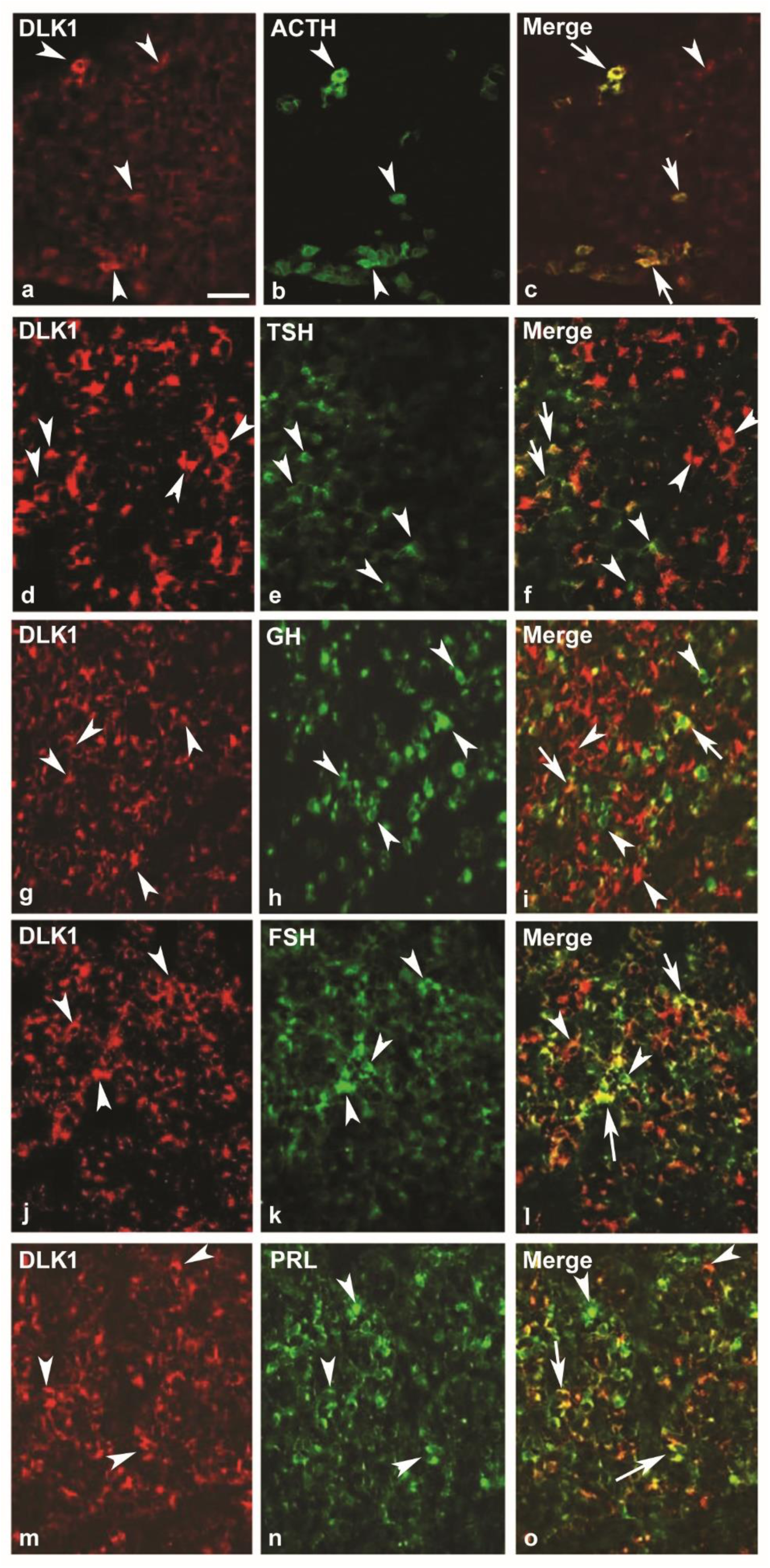
Double immunofluorescence showing DLK1 immunoreactivity in relation to hormone-producing cell populations at e18.5 in the anterior lobe. (a–c) DLK1 and ACTH immunoreactivity. (d–f) DLK1 and TSH immunoreactivity. (g–i) DLK1 and GH immunoreactivity. (j–l) DLK1 and FSH immunoreactivity. (m–o) DLK1 and PRL immunoreactivity. Merged images show double-labelled cells for DLK1 and pituitary hormones. Arrowheads indicate representative singly labelled cells, whereas arrows indicate representative double-labelled cells. Representative singly labelled DLK1-immunoreactive or hormone-immunoreactive cells are also indicated in the merged images to facilitate interpretation. Scale bar = 25 µm.

At this stage, DLK1-ir cells were detected throughout the IL and TL. Double-labelled cells for DLK1 and ACTH were observed in the IL (Figure 3), while double-labelled cells for DLK1 and FSH (Figure 4a–c), DLK1 and PRL (Figure 4d–f), and DLK1 and TSH (Figure 4g–i) were also observed in the TL. The distribution and relative abundance of double-labelled cells varied among endocrine cell populations and anatomical regions. Within the TL, qualitative examination suggested that DLK1 immunoreactivity appeared to be more frequently associated with PRL-immunoreactive cells than with FSH- or TSH-immunoreactive cells.

**Figure 3.**
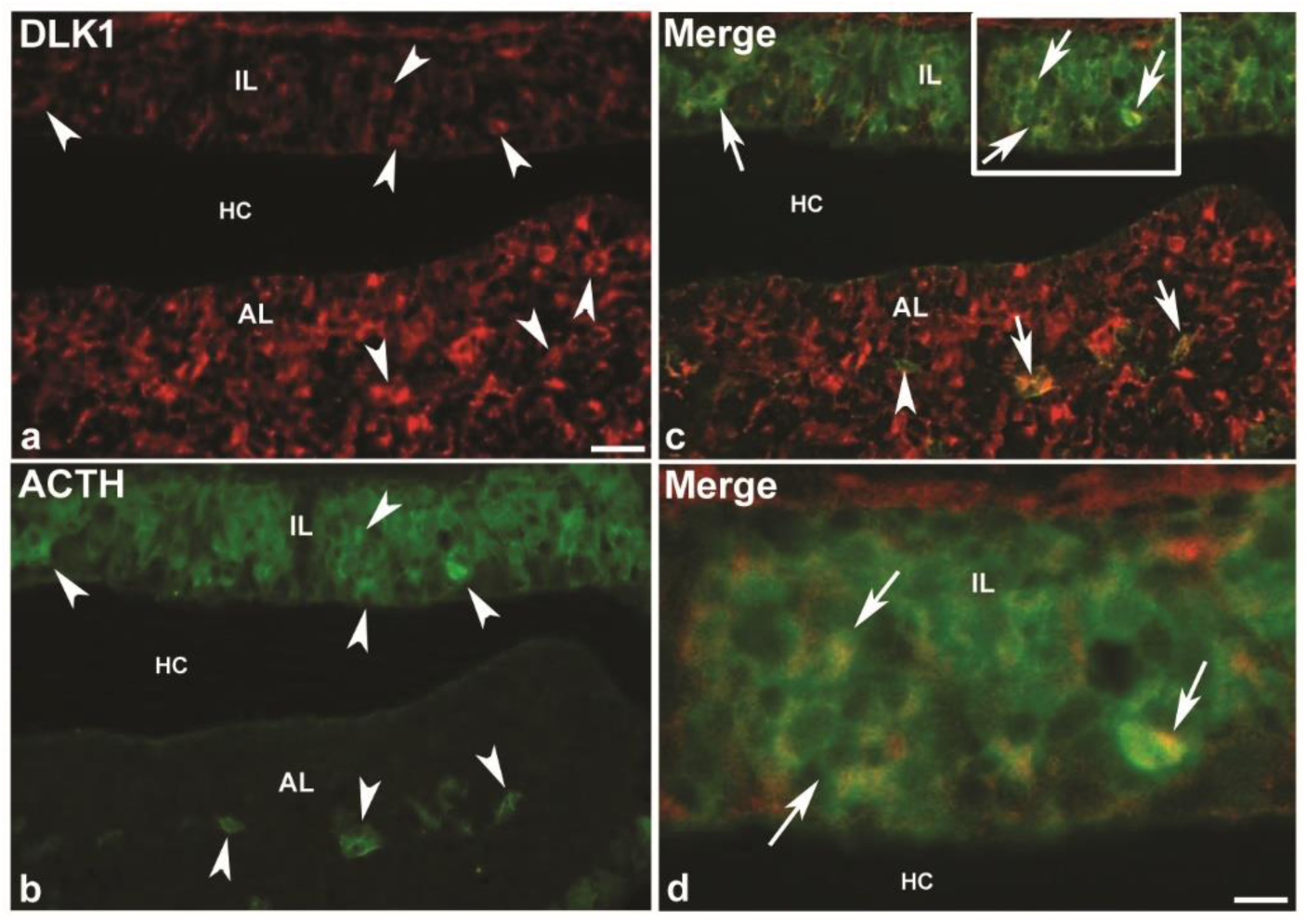
Double immunofluorescence showing DLK1 (a) and ACTH (b) immunoreactivity at e18.5 in the intermediate lobe. (c) Merged image showing double-labelled cells for DLK1 and ACTH. (d) Higher-magnification view of the boxed area in the intermediate lobe shown in (c). Arrowheads indicate representative singly labelled cells, whereas arrows indicate representative double-labelled cells. Representative singly labelled DLK1- or ACTH-immunoreactive cells are also indicated in the merged images to facilitate interpretation. AL, anterior lobe; HC, hypophysary cleft; IL, intermediate lobe. Scale bars = (a–c) 25 µm; (d) 10 µm.

**Figure 4.**
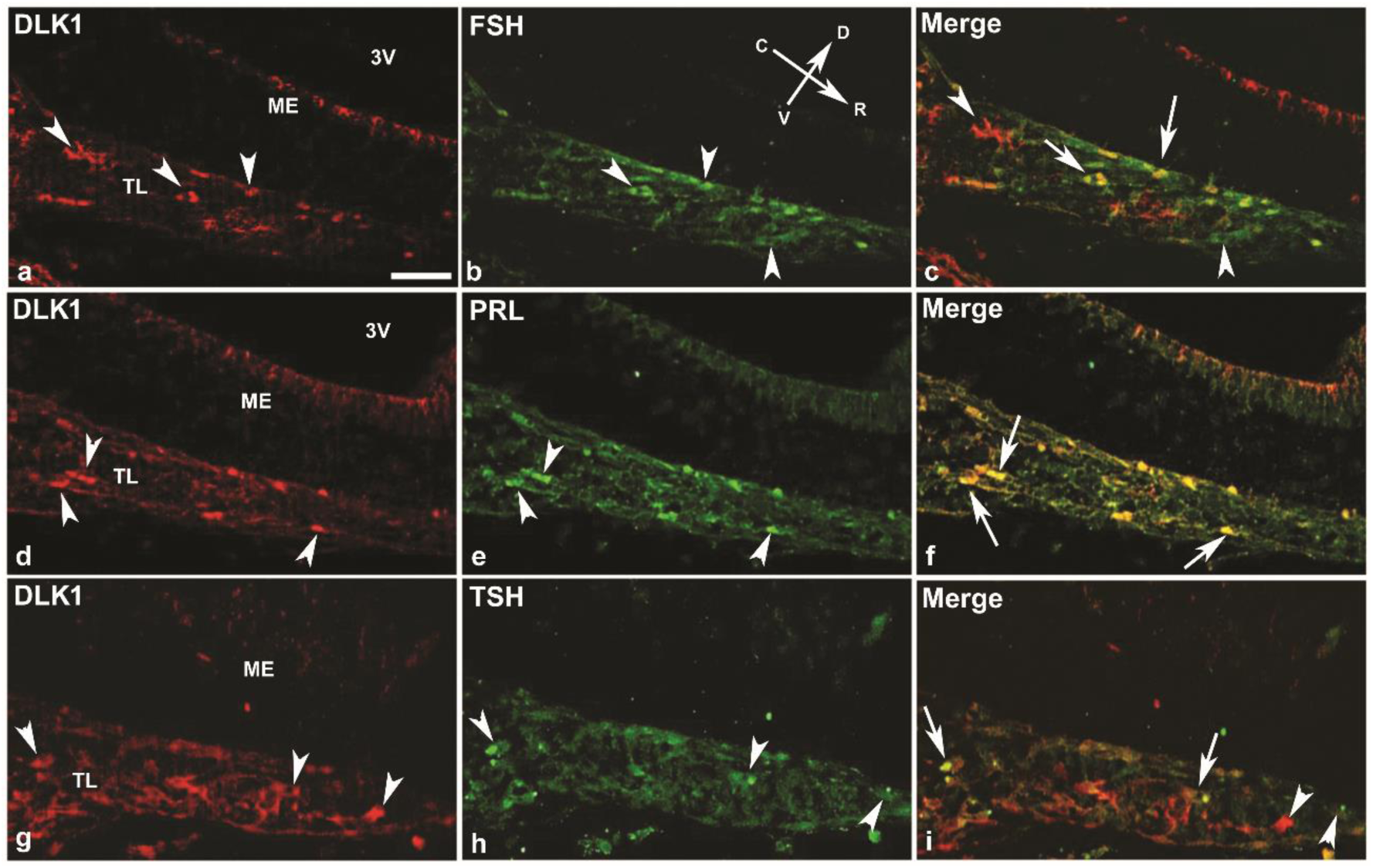
Double immunofluorescence showing DLK1 immunoreactivity in relation to hormone-producing cell populations at e18.5 in the tuberal lobe. (a-c) DLK1 and FSH immunoreactivity. (d-f) DLK1 and PRL immunoreactivity. (g-i) DLK1 and TSH immunoreactivity. Merged images (c,f,i) show representative double-labelled cells for DLK1 and pituitary hormones. Arrowheads indicate representative singly labelled cells, whereas arrows indicate representative double-labelled cells. Representative singly labelled DLK1-immunoreactive or hormone-immunoreactive cells are also indicated in the merged images to facilitate interpretation. ME, Median eminence; TL, Tuberal lobe. Scale bar = 25 µm.

### DLK1 immunoreactivity in postnatal pituitary hormone-producing cell populations

Double immunofluorescence analysis was performed at p15, when the major pituitary hormone-producing cell populations are differentiated. At p15, double-labelled cells for DLK1 and GH (Figure 5a–c), DLK1 and PRL (Figure 5d–f), DLK1 and ACTH (Figure 5g–i), DLK1 and TSH (Figure 5j–l) and DLK1 and FSH (Figure 5m–o) were identified in the AL. The proportion and distribution of double-labelled cells differed among the analyzed endocrine populations.

**Figure 5.**
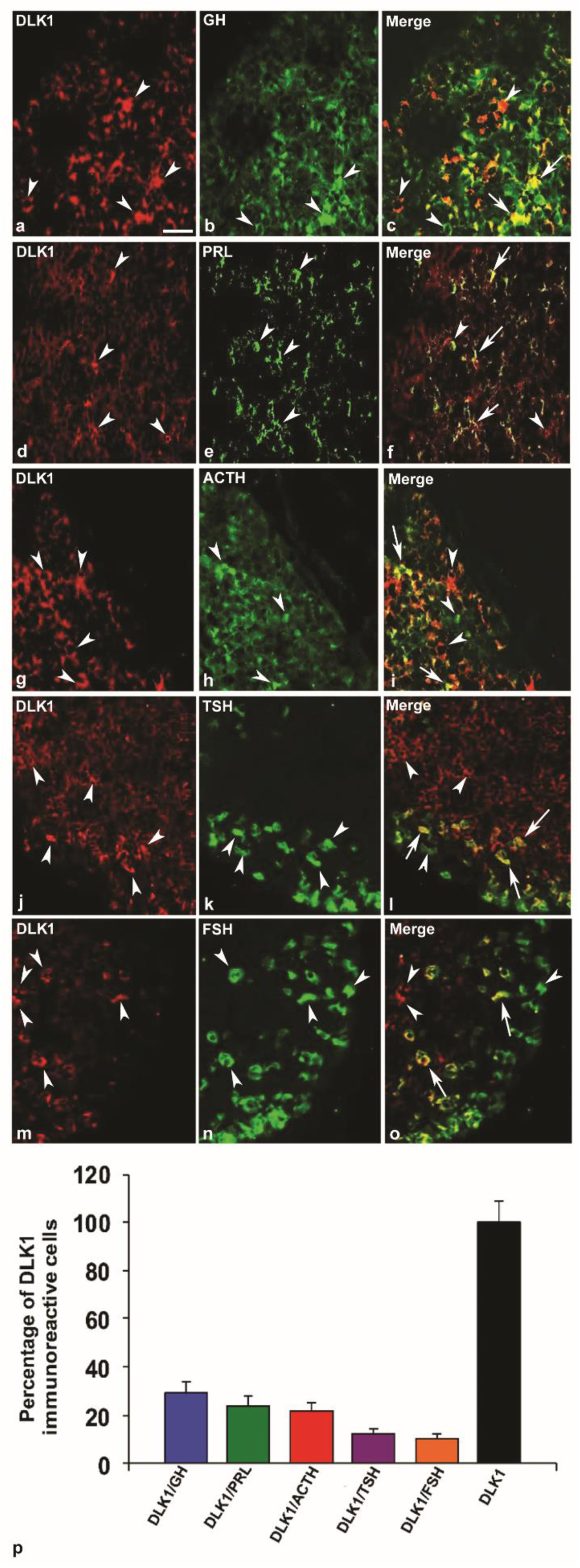
Double immunofluorescence and semiquantitative analysis of DLK1 immunoreactivity in relation to hormone-producing cell populations at p15 in the anterior pituitary lobe. (a–c) DLK1 and GH immunoreactivity. (d–f) DLK1 and PRL immunoreactivity. (g–i) DLK1 and ACTH immunoreactivity. (j–l) DLK1 and TSH immunoreactivity. (m–o) DLK1 and FSH immunoreactivity. Merged images illustrate representative examples of double-labelled cells for DLK1 and the corresponding pituitary hormone. Arrowheads indicate representative singly labelled cells, whereas arrows indicate representative double-labelled cells. Representative singly labelled DLK1-immunoreactive or hormone-immunoreactive cells are also indicated in the merged images to facilitate interpretation. (p) Semiquantitative estimation of the total number of DLK1-immunoreactive (DLK1-ir) cells in the anterior pituitary lobe and the percentage of DLK1-ir cells that were double-labelled for each pituitary hormone (n = 4). Values are expressed as mean ± SD. Scale bar = 25 µm.

### Semiquantitative analysis of DLK1-immunoreactive cells

Semiquantitative analysis of the anterior pituitary lobe at p15 estimated that approximately 32 ± 9% of adenohypophyseal cells displayed DLK1 immunoreactivity (Figure 5p). Semiquantitative analysis estimated that approximately 29 ± 5% of DLK1-ir cells were double-labelled for GH, 24 ± 4% with PRL, 22 ± 3% with ACTH, 12 ± 2% with TSH and 9 ± 2% with FSH (Figure 5p).

## Discussion

The present study describes the spatiotemporal distribution of DLK1 immunoreactivity during embryonic and postnatal development of the mouse pituitary gland, including its relationship with differentiating hormone-producing cell populations. DLK1-ir cells were detected from early stages of Rathke’s pouch formation and subsequently distributed throughout adenohypophyseal regions, the developing posterior pituitary and the median eminence during pituitary ontogeny. In addition, double-labelled cells for DLK1 and adenohypophyseal hormones were identified at e18.5 and p15.

Pituitary development is regulated by complex interactions among signalling pathways and transcription factors that coordinate stem/progenitor cells maintenance, regionalization and endocrine differentiation.^7,42,45^ Among these pathways, NOTCH signalling participates in maintenance of pituitary stem/progenitor cells populations and regulation of endocrine lineage differentiation.^17,18,43,44^ Canonical NOTCH pathway components, including receptors, ligands and downstream effectors, display dynamic spatial and temporal expression patterns during pituitary organogenesis.^17,18,43^

DLK1 is considered a non-canonical modulator of NOTCH signalling, although its precise mechanisms of action remain incompletely understood.^19,46^ Previous studies have demonstrated that DLK1 may interact with NOTCH receptors and modulate NOTCH signalling activity.^21–25^ In addition, DLK1 expression has been associated with progenitor and immature cell populations in several developmental systems, where it has been implicated in the regulation of cellular differentiation processes.^26–28^ In the pituitary gland, DLK1 expression has been previously associated with stem cell populations and endocrine differentiation processes.^33,42^ These observations partially parallel previous descriptions of canonical NOTCH pathway components during pituitary development,^18,17^ although the present data do not allow direct functional relationships between DLK1 and NOTCH signalling to be established.

Our observations are generally consistent with previous studies in rodents reporting Dlk1 mRNA or DLK1 protein expression in developing pituitary tissues.^32,33^ However, some regional and temporal differences appear to exist between mouse and rat pituitary development. In the mouse, DLK1 immunoreactivity progressively decreased in the intermediate lobe during postnatal stages, whereas DLK1-ir persisted in the anterior lobe, posterior pituitary and median eminence. In contrast, previous studies in rats reported earlier reduction of DLK1 immunoreactivity in neurohypophyseal regions.^32^ These interspecific differences may reflect species-dependent patterns of pituitary maturation and regional differentiation.

Double immunofluorescence analyses demonstrated the presence of double-labelled cells for DLK1 together with ACTH, GH, PRL, TSH and FSH at e18.5 and p15. Nevertheless, the distribution and relative abundance of double-labelled cells varied among pituitary regions and endocrine cell populations. In agreement with previous studies in rodents and humans, GH-producing cells represented one of the major endocrine populations associated with DLK1 immunoreactivity.^29,32,47^ Since GH-producing cells increase substantially during late embryonic and postnatal development,^48,49^ the relative proportions observed in the present semiquantitative analyses likely reflect the developmental stage examined.

Nevertheless, the coexistence observed between DLK1 and the different pituitary hormones indicates a spatial association between DLK1-immunoreactive cells and differentiating endocrine cell populations during pituitary development. This association was observed across several endocrine populations, although its relative extent varied according to the hormone and developmental stage examined.

The widespread distribution of DLK1 during early stages of Rathke’s pouch development, together with its subsequent persistence in differentiating endocrine populations, is compatible with a potential association between DLK1 expression and transitional states of pituitary cell differentiation. However, the present study is descriptive in nature and does not establish functional roles for DLK1 during pituitary organogenesis or endocrine differentiation.

An additional finding of this study was the persistence of DLK1 immunoreactivity in both the median eminence and the posterior pituitary during postnatal stages. DLK1-ir fibers were observed in the median eminence, whereas DLK1-ir persisted within the posterior lobe. Previous studies have demonstrated the involvement of NOTCH pathway effectors in neurohypophyseal development and pituicyte differentiation.^50,51^ Although the relationship between DLK1 and these processes remains unclear, the present observations support the possibility that DLK1 expression is maintained in both the adenohypophysis and the posterior pituitary, as well as in the median eminence, during pituitary maturation. In conclusion, the present study provides a detailed description of DLK1 immunoreactivity during mouse pituitary ontogeny and documents dynamic spatial and temporal distribution patterns in the adenohypophysis, posterior pituitary and median eminence. Furthermore, DLK1 immunoreactivity was observed in association with differentiating hormone-producing cell populations during embryonic and postnatal development. However, further functional studies will be necessary to clarify the role of DLK1 during pituitary development and adult pituitary physiology.

## Availability of data and materials

All data generated or analysed during this study are included in this manuscript and its Supplementary Material. Further inquiries can be directed to the corresponding author.

## Acknowledgements

The authors are very grateful to Dr Gerard Tramu (University of Bordeaux 1, Bordeaux, France) for providing hormone antisera against ACTH, TSH and FSH.

## Declaration of generative artificial intelligence and artificial intelligence-assisted technologies in the writing process

During the preparation of this manuscript, the authors used ChatGPT (OpenAI) to assist with language editing and improving the clarity of the text. The authors reviewed and edited the content as needed and take full responsibility for the content of the manuscript.

---

**Figure S1.**
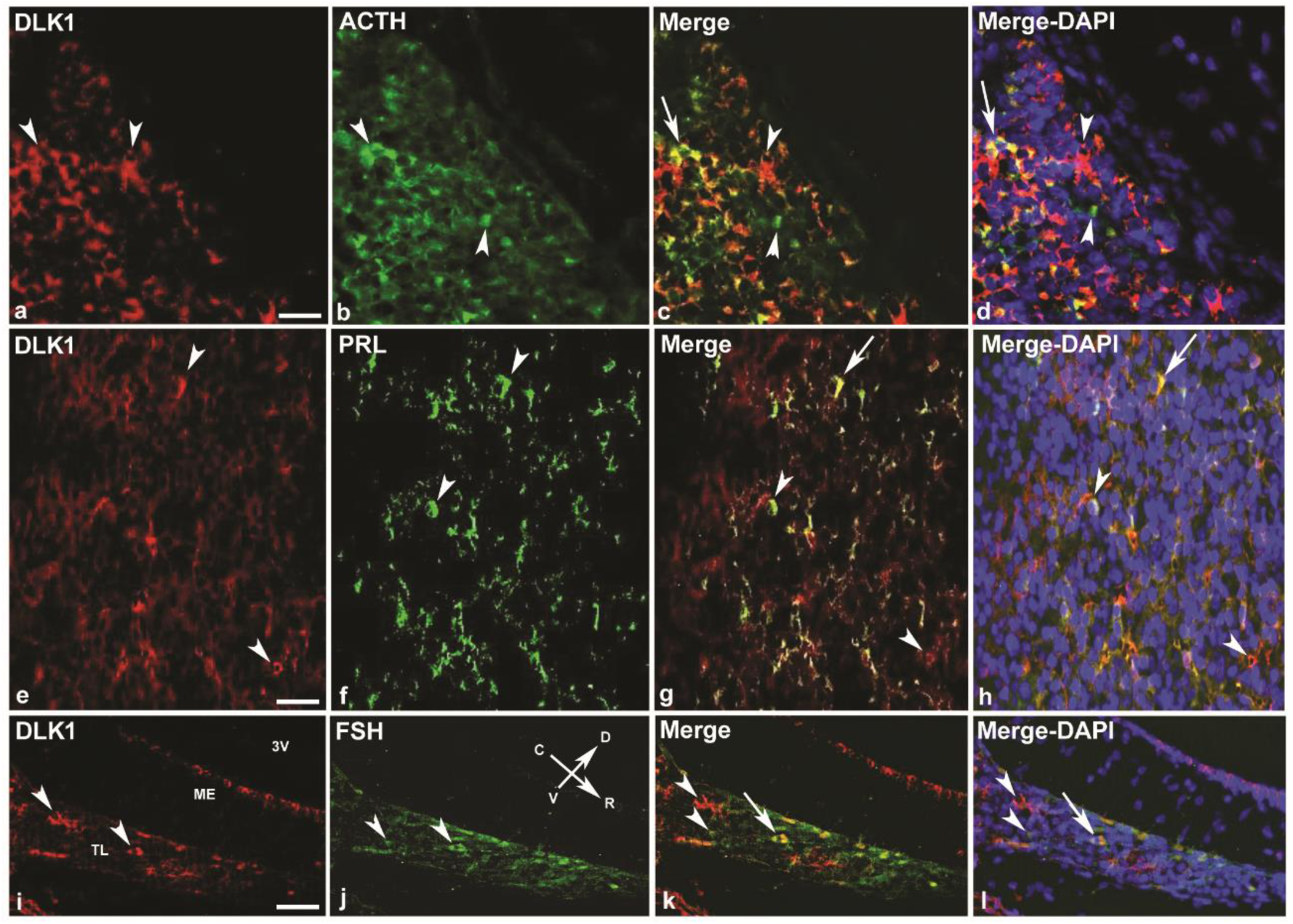
Representative examples illustrating DAPI nuclear counterstaining in double immunofluorescence preparations of the developing mouse pituitary. (a–d) DLK1 and ACTH immunoreactivity in the anterior pituitary lobe at postnatal day 15 (p15). (e–h) DLK1 and PRL immunoreactivity in the anterior pituitary lobe at p15. (i–l) DLK1 and FSH immunoreactivity in the tuberal lobe at embryonic day 18.5 (e18.5). In each series, the first and second panels show DLK1 and hormone immunoreactivity, respectively, followed by merged images without (third column) and with DAPI nuclear counterstaining (fourth column). The merged images including DAPI demonstrate preserved tissue integrity and high cellular density of the adenohypophyseal tissue, while also illustrating that the intense DAPI nuclear signal partially masks the comparatively weaker DLK1 and hormone immunofluorescent signals, thereby reducing the visual clarity of double-labelled cells. For this reason, merged images presented in the main figures were generated without DAPI counterstaining to facilitate visualization of DLK1-hormone immunoreactive cells. Arrowheads indicate representative singly labelled cells, whereas arrows indicate representative double-labelled cells. Scale bars = 25 µm.

**Figure S2.**
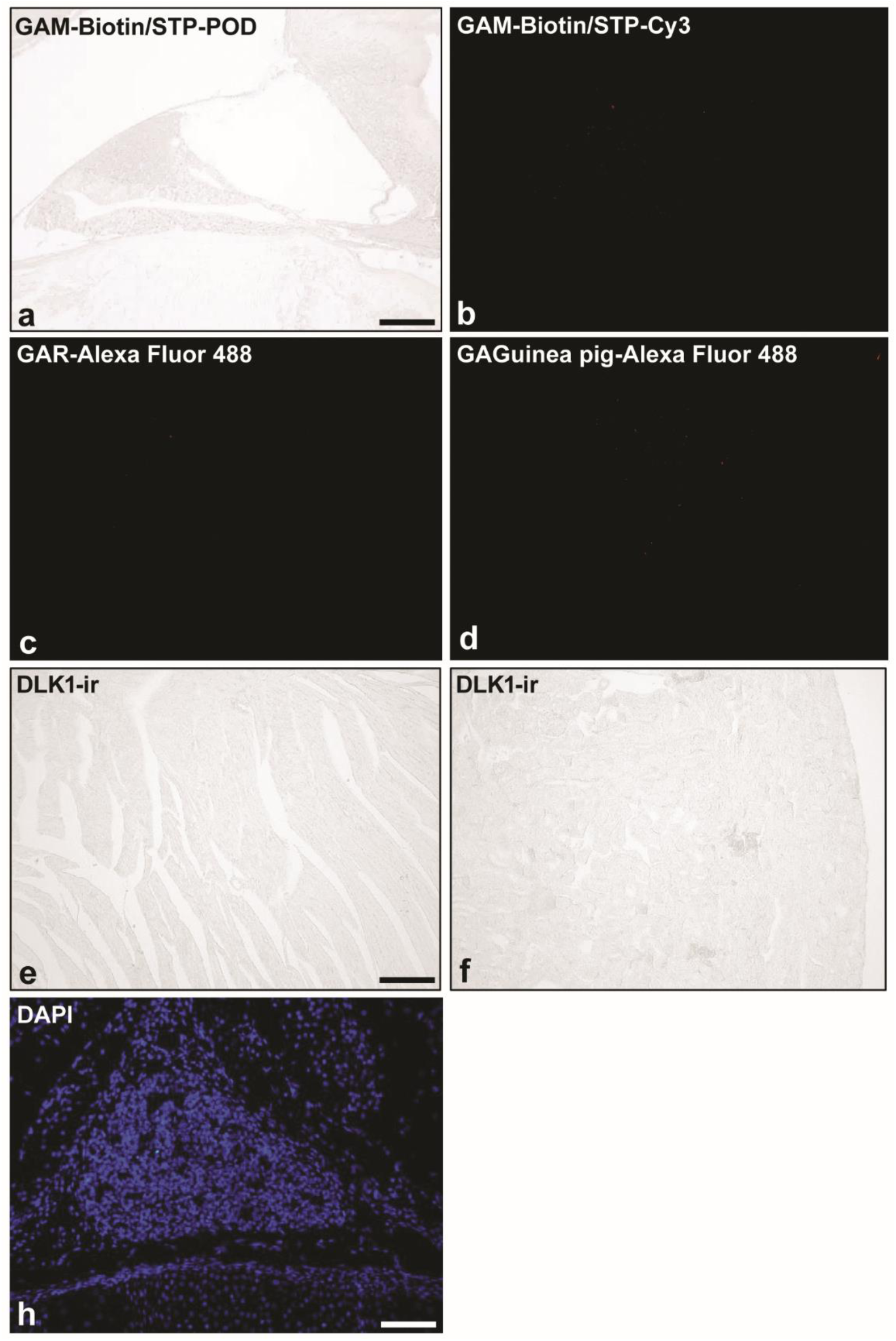
Negative control sections processed under identical conditions, either by omission of the corresponding primary antibodies or using tissues reported to lack detectable DLK1 immunoreactivity. No specific signal was detected in either the chromogenic or fluorescence procedures. For DLK1 immunohistochemistry, the mouse-specific secondary antibody was followed by POD-conjugated streptavidin (a), whereas for DLK1 immunofluorescence, the same secondary antibody was followed by Cy3-conjugated streptavidin (b). ACTH, TSH, FSH and GH antisera were detected using the same rabbit-specific secondary antibody conjugated to Alexa Fluor 488 (c), whereas PRL was detected using a guinea pig-specific secondary antibody conjugated to Alexa Fluor 488 (d). Therefore, representative negative controls for these antisera are shown. Adult mouse heart and kidney sections were also processed for DLK1 immunohistochemistry using the immunoperoxidase procedure described by Shu et al. (1988) and showed no specific immunoreactivity (e, f). (g) Representative DAPI-stained sagittal section of the e18.5 pituitary gland showing preserved tissue integrity and morphology. Scale bars = 60 µm.

## References

1. Batista MA, Doerr-Schott J, Bello AR. Immunohistochemical study on the development of the adenohypophyseal cells in the lizard Gallotia galloti. Anat Embryol 1989;180:143–9.

2. Chapman SC, Sawitzke AL, Campbell DS, Schoenwolf GC. A three-dimensional atlas of pituitary gland development in the zebrafish. J Comp Neurol 2005;487:428–40.

3. Reyes R, Martínez S, González M, Tramu G, Bello AR. Origin of adenohypophyseal lobes and cells from Rathke’s pouch in Swiss albino mice. Proliferation and expression of Pitx2 and Calbindin D28K in corticotropic and somatotropic cell differentiation. Anat Histol Embryol 2008;37:263–71.

4. Reyes R, González M, Bello AR. Origin of adenohypophyseal lobes and cells from Rathke’s pouch in chicken (Gallus gallus) and Japanese quail (Coturnix coturnix japonica). Expression of calcium-binding proteins. Anat Histol Embryol 2008;37:272–82.

5. Sánchez-Arrones L, Ferrán JL, Hidalgo-Sanchez M, Puelles L. Origin and early development of the chicken adenohypophysis. Front Neuroanat 2015;9:7.

6. Scagliotti V, Costa Fernandes Esse R, Willis TL, Howard M, Carrus I, Lodge E, et al. Dynamic expression of imprinted genes in the developing and postnatal pituitary gland. Genes 2021;12:509.

7. Edwards W, Raetzman LT. Complex integration of intrinsic and peripheral signaling is required for pituitary gland development. Biol Reprod 2018;99:504–13.

8. Rizzoti K. Genetic regulation of murine pituitary development. J Mol Endocrinol 2015;54:R55–R73.

9. Kelberman D, Rizzoti K, Lovell-Badge R, Robinson IC, Dattani MT. Genetic regulation of pituitary gland development in human and mouse. Endocr Rev 2009;30:790–829.

10. Zhu X, Gleiberman AS, Rosenfeld MG. Molecular physiology of pituitary development: Signaling and transcriptional networks. Physiol Rev 2007;87:933–63.

11. Zhu X, Wang J, Ju BG, Rosenfeld MG. Signaling and epigenetic regulation of pituitary development. Curr Opin Cell Biol 2007;19:605–11.

12. Quentien MH, Barlier A, Franc JL, Pellegrini I, Brue T, Enjalbert A. Pituitary transcription factors: From congenital deficiencies to gene therapy. J Neuroendocrinol 2006;18:633–42.

13. Dasen JS, Rosenfeld MG. Signaling and transcriptional mechanisms in pituitary development. Annu Rev Neurosci 2001;24:327–55.

14. Parkinson N, Collins MM, Dufresne L, Ryan AK. Expression patterns of hormones, signaling molecules, and transcription factors during adenohypophysis development in the chick embryo. Dev Dyn 2010;239:1197–210.

15. Sornson MW, Wu W, Dasen JS, Flynn SE, Norman DJ, O’Connell SM, et al. Pituitary lineage determination by the Prophet of Pit-1 homeodomain factor defective in Ames dwarfism. Nature 1996;384:327–33.

16. Dutta S, Dietrich JE, Westerfield M, Varga ZM. Notch signaling regulates endocrine cell specification in the zebrafish anterior pituitary. Dev Biol 2008;319:248–57.

17. Zhu X, Zhang J, Tollkuhn J, Ohsawa R, Bresnick EH, Guillemot F, et al. Sustained Notch signaling in progenitors is required for sequential emergence of distinct cell lineages during organogenesis. Genes Dev 2006;20:2739–53.

18. Raetzman LT, Ross SA, Cook S, Dunwoodie SL, Camper SA, Thomas PQ. Developmental regulation of Notch signaling genes in the embryonic pituitary: Prop1 deficiency affects Notch2 expression. Dev Biol 2004;265:329–40.

19. D’Souza B, Miyamoto A, Weinmaster G. The many facets of Notch ligands. Oncogene 2008;27:5148–67.

20. Mumm JS, Kopan R. Notch signaling: From the outside in. Dev Biol 2000;228:151–65.

21. Traustadóttir G, Jensen CH, Thomassen M, Beck HC, Mortensen SB, Laborda J, et al. Evidence of non-canonical NOTCH signaling: Delta-like 1 homolog (DLK1) directly interacts with the NOTCH1 receptor in mammals. Cell Signal 2016;28:246–54.

22. Traustadóttir G, Jensen CH, Garcia Ramirez JJ, Beck HC, Sheikh SP, Andersen DC. The non-canonical NOTCH1 ligand Delta-like 1 homolog (DLK1) self interacts in mammals. Int J Biol Macromol 2017;97:460–7.

23. Bray SJ, Takada S, Harrison E, Shen SC, Ferguson-Smith AC. The atypical mammalian ligand Delta-like homologue 1 (Dlk1) can regulate Notch signalling in Drosophila. BMC Dev Biol 2008;8:11.

24. Nueda ML, Baladrón V, Sánchez-Solana B, Ballesteros MA, Laborda J. The EGF-like protein Dlk1 inhibits Notch signaling and potentiates adipogenesis of mesenchymal cells. J Mol Biol 2007;367:1281–93.

25. Baladrón V, Ruiz-Hidalgo MJ, Nueda ML, Díaz-Guerra MJ, García-Ramírez JJ, Bonvini E, et al. Dlk acts as a negative regulator of Notch1 activation through interactions with specific EGF-like repeats. Exp Cell Res 2005;303:343–59.

26. Mirshekar-Syahkal B, Haak E, Kimber GM, van Leusden K, Harvey K, O’Rourke J, et al. Dlk1 is a negative regulator of emerging hematopoietic stem and progenitor cells. Haematologica 2013;98:163–71.

27. Garcés C, Ruiz-Hidalgo MJ, Bonvini E, Goldstein J, Laborda J. Adipocyte differentiation is modulated by secreted delta-like (dlk) variants and requires the expression of membrane-associated dlk. Differentiation 1999;64:103–14.

28. Smas CM, Sul HS. Pref-1, a protein containing EGF-like repeats, inhibits adipocyte differentiation. Cell 1993;73:725–34.

29. Ansell PJ, Zhou Y, Schjeide BM, Kerner A, Zhao J, Zhang X, Klibanski A. Regulation of growth hormone expression by Delta-like protein 1 (Dlk1). Mol Cell Endocrinol 2007;271:55–63.

30. Puertas-Avendaño RA, González-Gómez MJ, Ruvira MD, Ruiz-Hidalgo MJ, Morales-Delgado N, Laborda J, et al. Role of the non-canonical Notch ligand Delta-like protein 1 in hormone-producing cells of the adult male mouse pituitary. J Neuroendocrinol 2011;23:849–59.

31. Jensen CH, Meyer M, Schroder HD, Kliem A, Zimmer J, Teisner B. Neurons in the monoaminergic nuclei of the rat and human central nervous system express FA1/Dlk. Neuroreport 2001;12:3959–63.

32. Nakakura T, Sato M, Suzuki M, Hatano O, Takemori H, Taniguchi Y, et al. The spatial and temporal expression of delta-like protein 1 in the rat pituitary gland during development. Histochem Cell Biol 2009;131:141–53.

33. Scagliotti V, Vignola ML, Willis T, Howard M, Marinelli E, Gaston-Massuet C, et al. Imprinted Dlk1 dosage as a size determinant of the mammalian pituitary gland. eLife 2023;12:e84092.

34. Bard JBL, Kaufman MH, Dubreuil C, Brune RM, Burger A, Baldock RA, Davidson DR. An internet-accessible database of mouse developmental anatomy based on a systematic nomenclature. Mech Dev 1998;74:111–20.

35. Shu SY, Ju G, Fan LZ. The glucose oxidase-DAB-nickel method in peroxidase histochemistry of the nervous system. Neurosci Lett 1988;85:169–71.

36. Dubois MP. Localisation par immunocytologie des hormones glycoprotidiques hypophysaires chez le vertébrés. In: Les hormones glycoprotéiques hypophysaires. Colloque INSERM, Hôpital Saint Antoine. Paris: Inserm; 1972a. pp 27–47.

37. Dubois P. Localisation cytologique par immunofluorescence des sécrétions corticotropes, α et β-mélanotropes au niveau de l’antéhypophyse des bovins, ovins et porcins. Z Zellforsch Mikrosk Anat 1972b;125:200–9.

38. Tramu G, Dubois MP. Comparative cellular localisation of corticotropin and melanotropin in lerot adenohypophysis (Eliomys quercinus). An immunohistochemical study. Cell Tissue Res 1977;183:457–69.

39. Bello AR, Dubourg P, Kah O, Tramu G. Identification of the neurotensin immunoreactive cells in the anterior pituitary of normal and castrated rats: a double immunocytochemical investigation at the light and electron microscope levels. Neuroendocrinology 1992;55:714–22.

40. Falix FA, Tjon-A-Loi MRS, Gaemers IC, Aronson DC, Lamers WH. DLK1 protein expression during mouse development provides new insights into its function. ISRN Dev Biol 2013;2013:628962.

41. Van de Pette M, Domond A, Galvão AM, Millership SJ, To W, Prodani Ch, et al. Epigenetic changes induced by in utero dietary challenge result in phenotypic variability in successive generations of mice. Nat Commun 2022;13:2464.

42. Cheung LYM, Rizzoti K, Lovell-Badge R, Le Tissier PR. Pituitary phenotypes of mice lacking the Notch signalling ligand Delta-like 1 homologue. J Neuroendocrinol 2013;25:391–401.

43. Kita A, Imayoshi I, Hojo M, Kitagawa M, Kokubu H, Ohsawa R, et al. Hes1 and Hes5 control the progenitor pool, intermediate lobe specification, and posterior lobe formation in pituitary development. Mol Endocrinol 2007;21:1458–66.

44. Nantie LB, Himes AD, Getz DR, Raetzman LT. Notch signaling in postnatal pituitary expansion: Proliferation, progenitors, and cell specification. Mol Endocrinol 2014;28:731–44.

45. Haston S, Manshaei S, Martinez-Barbera JP. Stem/progenitor cells in pituitary organ homeostasis and tumourigenesis. J Endocrinol 2018;236:R1–R13.

46. Falix FA, Aronson DC, Lamers WH, Gaemers IC. Possible roles of DLK1 in the Notch pathway during development and disease. Biochim Biophys Acta 2012;1822:988–95.

47. Bello AR, Puertas-Avendaño RA, González-Gómez MJ, González-Gómez M, Laborda J, Damas C, et al. Delta-like protein 1 in the pituitary-adipose axis in the adult male mouse. J Neuroendocrinol 2017;29:e12507.

48. Bjelobaba I, Janjic MM, Kucka M, Stojilkovic SS. Cell type-specific sexual dimorphism in rat pituitary gene expression during maturation. Biol Reprod 2015;93:21.

49. Simmons DM, Voss JW, Ingraham HA, Holloway JM, Broide RS, Rosenfeld MG, Swanson LW. Pituitary cell phenotypes involve cell-specific Pit-1 mRNA translation and synergistic interactions with other classes of transcription factors. Genes Dev 1990;4:695–711.

50. Aujla PK, Bora A, Monahan P, Sweedler JV, Raetzman LT. The Notch effector gene Hes1 regulates migration of hypothalamic neurons, neuropeptide content and axon targeting to the pituitary. Dev Biol 2011;353:61–71.

51. Goto M, Hojo M, Ando M, Kita A, Kitagawa M, Ohtsuka T, et al. Hes1 and Hes5 are required for differentiation of pituicytes and formation of the neurohypophysis in pituitary development. Brain Res 2015;1625:206–17.

